# The guidance and adhesion protein FLRT2 dimerizes *in cis* via dual Small-X_3_-Small transmembrane motifs

**DOI:** 10.1101/2020.10.06.328401

**Authors:** Verity Jackson, Julia Hermann, Christopher J. Tynan, Daniel J. Rolfe, Robin A. Corey, Anna L. Duncan, Maxime Noriega, Amy Chu, Antreas C. Kalli, E. Yvonne Jones, Mark S. P. Sansom, Marisa L. Martin-Fernandez, Elena Seiradake, Matthieu Chavent

## Abstract

Fibronectin Leucine-rich Repeat Transmembrane (FLRT 1-3) proteins are a family of broadly expressed single-spanning transmembrane receptors that play key roles in development. Their extracellular domains mediate homotypic cell-cell adhesion and heterotypic protein interactions with other receptors to regulate cell adhesion and guidance. These *in trans* FLRT interactions determine the formation of signaling complexes of varying complexity and function. Whether FLRTs also interact at the surface of the same cell, *in cis*, remains unknown. Here, molecular dynamics simulations reveal two dimerization motifs in the FLRT2 transmembrane helix. Single particle tracking experiments show that these ‘Small-X_3_-Small’ motifs synergize with a third dimerization motif encoded in the extracellular domain to permit the *cis* association and co-diffusion patterns of FLRT2 receptors on cells. These results may point to a competitive switching mechanism between *in cis* and *in trans* interactions which suggests that homotypic FLRT interaction mirrors the functionalities of classic adhesion molecules.

**Fields:** Structural Biology and Biophysics / Computational Biology

## Introduction

Fibronectin Leucine-rich Repeat Transmembrane (FLRT) proteins are a family of cell adhesion molecules (CAMs) that are broadly expressed during vertebrate development (Karaulanov et al., 2006; Maretto et al., 2008). FLRTs are unusual CAMs as they perform both cell adhesive and repulsive functions, leading to their definition as Repellent CAMs (ReCAMs) (Seiradake et al., 2014; Yamagishi et al., 2011). In neurons, FLRTs act as repulsive guidance cues during cortical cell migration (Jackson et al., 2015; Yamagishi et al., 2011), where they play a key role in cortical folding (Toro et al., 2017) and as adhesion molecules in synaptic complexes (O’Sullivan et al., 2012; Sando et al., 2019). Adhesive FLRT functions are elicited by homotypic binding (Karaulanov et al., 2006; Maretto et al., 2008) or by binding to the G-protein coupled receptor Latrophilin (Lphn 1-3) (Jackson et al., 2015; Lu et al., 2015; O’Sullivan et al., 2012; Ranaivoson et al., 2015) on opposing cells, while cell repulsion results from interaction with Uncoordinated-5 (Unc5A-D) (Lu et al., 2015; Yamagishi et al., 2011). FLRT also interacts with Unc5 *in cis* to regulate Lphn-mediated adhesion, at least *in vitro* (Jackson et al., 2016). In migrating neurons, FLRT cooperates with the Lphn-binding receptor Teneurin to form a ternary trans-synaptic complex that mediates cell repulsion (Toro et al., 2020), while the three proteins also function in promoting synapsing (Sando et al., 2019). Thus, FLRT acts in a context-dependent manner to determine the formation of different higher order cell-guidance signaling complexes and regulate brain development (Seiradake et al., 2016). Here we ask whether FLRT forms homotypic *cis* complexes and how this may modulate *cis* and *trans* interactions with other partners.

FLRTs share a common architecture (**Fig. 1A**) beginning with an N-terminal Leucine-Rich Repeat (LRR) extracellular domain, which contains a concave surface on which both FLRT and Lphn bind (Jackson et al., 2015; Seiradake et al., 2014). Unc5 binds to an adjacent surface on the LRR domain, which is compatible at least with Lphn-binding (Jackson et al., 2016). The LRR domain is linked to a type III fibronectin (FN) domain which then leads into the single-spanning transmembrane (TM) domain and a ~100 amino acid long intracellular domain (ICD) of unknown structure. FLRT2 TM domains contain two consecutives “Small-X_3_-Small” motifs (**Fig. 1B**) which are known to promote receptor interactions *in cis* (Russ and Engelman, 2000; Teese and Langosch, 2015). For example, this motif plays fundamental roles in the signaling mechanisms of epidermal growth factor receptor (EGFR), fibroblast growth factor receptor (FGFR), and EphA receptors (Bocharov et al., 2008; Endres et al., 2013; Sarabipour and Hristova, 2016).

**Fig. 1:**
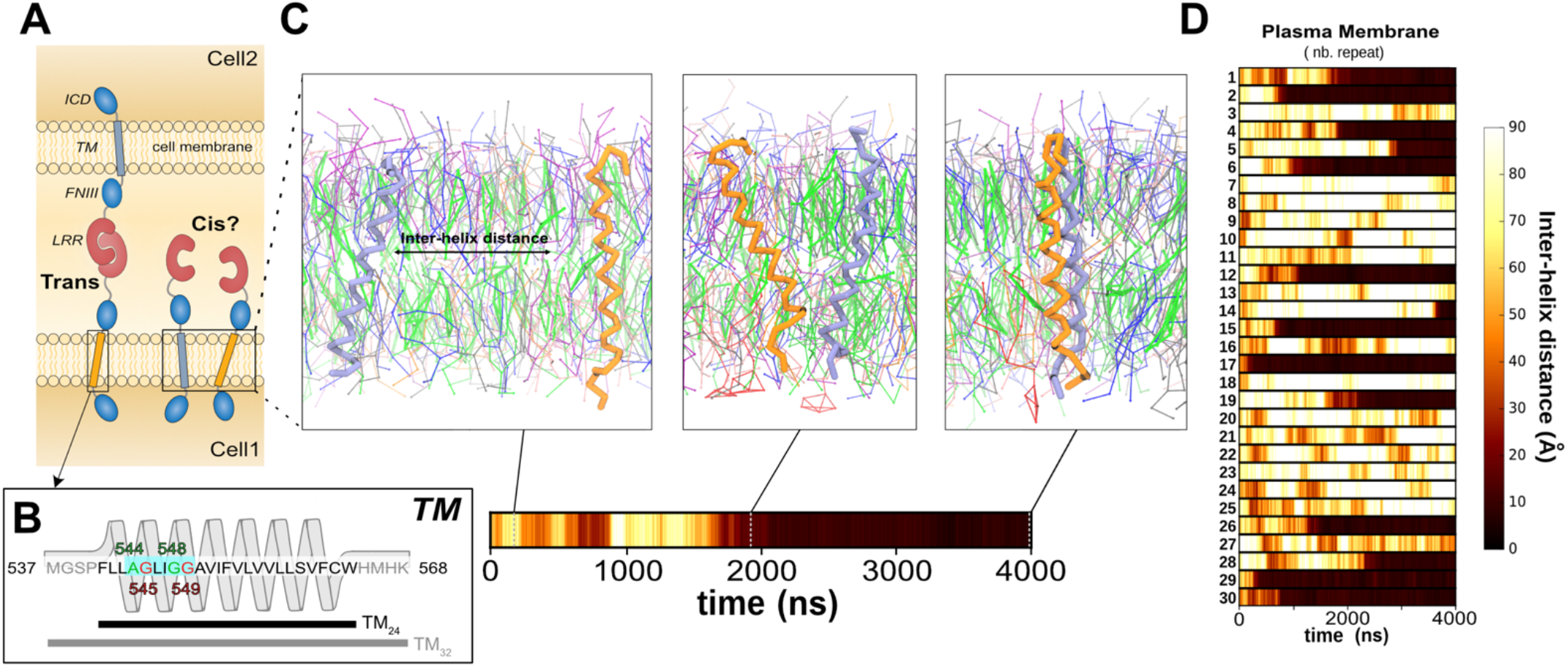
Dimerization of FLRT2 TM domains in the plasma membrane. **A-** Schematic of FLRT proteins engaging *in trans* and potentially also *in cis* interactions. **B-** Sequence of the FLRT2 TM helix. The two Small-X_3_-Small motifs, key residues for the formation of the helix/helix interface highlighted by the CG-MD simulations, are colored in green and red. Two constructs were used as inputs for MD simulations: the core TM helix of 24 residues (TM_24_) and an extended version with the four most N- and C-terminal residues (TM_32_). **C-** The CG-MD protocol to assess TM helix interactions. The two helices are positioned 60 Å apart and diffuse freely in the membrane. The colored bars show, for each simulation, the distance between the two TM helices as a function of time. **D-** The TM contact bars of the 30 simulations for the TM_32_ helices in the plasma membrane constituted of 8 different lipid types (see details in Sup. Fig. 1).

Characterizing the dynamics of membrane protein structure is challenging (Bugge et al., 2016), especially due to the interactions between lipids and proteins (Cymer et al., 2012; Laganowsky et al., 2014; Pliotas et al., 2015; Sonntag et al., 2011). Molecular Dynamics (MD) simulations have recently emerged as powerful tools to study membrane protein interactions (Chavent et al., 2016). In particular, coarse-grained (CG) modelling is a method of choice to explore the association of TM domains (Souza et al., 2021; Wassenaar et al., 2015a) in biological membranes (Corradi et al., 2018; Marrink et al., 2019). Combining MD simulations with experimental assays is now a well-established scientific strategy (Bottaro and Lindorff-Larsen, 2018). Conversely, Single Molecule Tracking (SMT) microscopy (Liu et al., 2016; Stone et al., 2017) provides the resolution and dynamic insight to validate models of the assembly mechanisms of cell receptors (Wilmes et al., 2020; Zanetti-Domingues et al., 2018).

Here, we use molecular dynamics simulations and live cell SMT experiments to reveal how FLRT2 dimerizes *in cis* via two Small-X_3_-Small motifs. Unexpectedly, these motifs work synergistically with the extracellular dimerization motif in the ligand-binding domain (Seiradake et al., 2014) to produce FLRT-FLRT association. The results suggest a bipartite structural mechanism that underlies the diverse functions of FLRT, and a competitive mechanism for *in cis* versus *in trans* binding via the extracellular domain.

## Results

### FLRT2 TM dimerization involves two Small-X_3_-Small motifs

As no structural information exists for the FLRT2 TM domain, we have used secondary structure prediction tools (see Methods) to predict the membrane-embedded helical region of FLRT2 (**Fig. 1B**). We identified 24 residues as the core TM helix (denoted TM_24_). This length is consistent with the average length for a plasma membrane-spanning TM helix (Sharpe et al., 2010). We extended the helical segment with four N- and C-terminal residues, which were modelled as coils (denoted TM_32_).

We performed multiple runs of coarse-grained molecular dynamics (CG-MD) (Marrink et al., 2007; Monticelli et al., 2008) to model the associations of the TM_32_ monomers in a membrane model composed of 8 different species of lipids (**Sup. Fig. 1**) mimicking to some extent the complexity of an average plasma membrane (PM) (Ingólfsson et al., 2020). We positioned the two helices 60 Å apart allowing them to diffuse freely until there is an encounter which mainly leads to the formation of a stable helix dimer (**Fig. 1C,D**). The helices interacted through a network of residues distributed along each peptide. Among these residues, we identified two consecutive Small-X_3_-Small motifs known to favor TM interactions (Russ and Engelman, 2000; Teese and Langosch, 2015): A_544_-X_3_-G_548_ and G_545_-X_3_-G_549_ (**Fig. 2A**).

**Fig. 2:**
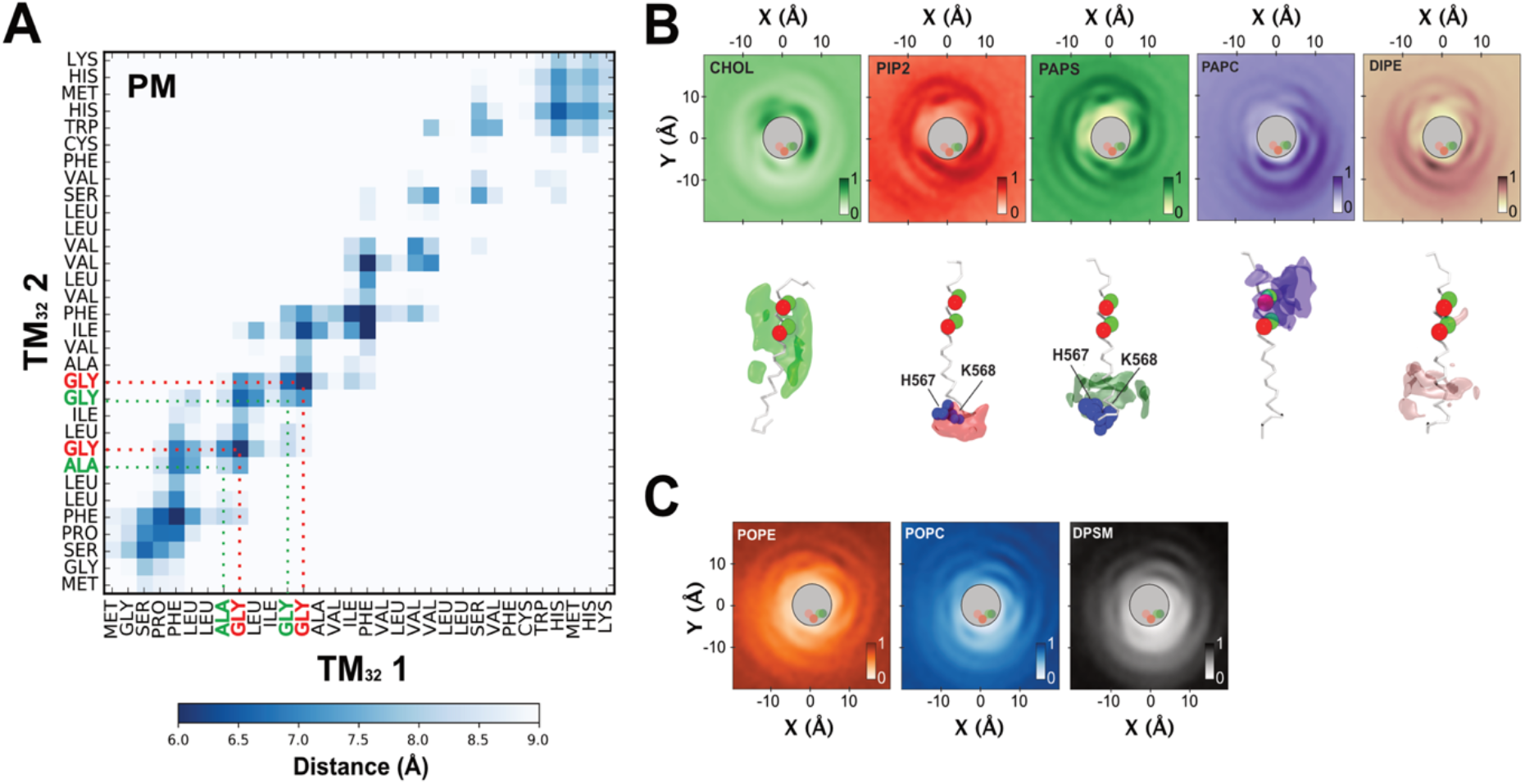
FLRT TM dimer interface and lipid fingerprint in the plasma membrane. **A-** Averaged TM contact matrix extracted from simulations of TM_32_ in plasma membrane (PM) highlighted a TM dimerization via the A_544_-X_3_-G_548_ and G_545_-X_3_-G_549_ motifs. Two-dimensional lateral density maps, showing local lipid density around one TM domain highlighting favored (**B**) and depleted lipids (**C**). For the lipids in the TM vicinity, a three dimensional representation of the lipid density displays a lipid redistribution spread along the whole TM domain.

Based on these simulations, we analyzed the composition of the lipid shell around a TM domain. This revealed preferential association with specific lipids (**Fig. 2B**): cholesterol, negatively charged lipids (PIP2 and PS), and highly unsaturated lipids (PAPC: C16:0/20:4, DIPE C16:2/C18:2). Due to the membrane asymmetry, these interactions were spread along the whole TM domain. Conversely, less saturated lipids (POPE and POPC) and sphingomyelin lipid (DPSM) seemed to be depleted from the direct surrounding of the TM domain (**Fig. 2C**). The interactions between the TM domain and surrounding lipids may create a unique membrane environment (Corradi et al., 2018) which accordingly may influence the dynamics of the TM dimerization.

### FLRT2 TM dimerization is a dynamical process modulated by membrane lipid composition

To assess the role of the membrane composition on the TM dimer dynamics we have defined three types of symmetric membrane: a membrane composed of unsaturated lipids (POPC: C16:0/18:1), a membrane composed of saturated lipids (DPPC: C16:0/18:0), and a lipid mixture of 80% DPPC and 20% cholesterol. For each composition, we ran multiple runs of coarse-grained molecular dynamics (CG-MD) simulations (**Sup. Table 1**). For the three membrane compositions, we mainly observed a dimerization of TM domains (**Sup. Fig. 2**).

We then performed crossing angle analysis to assess the geometry of the TM helices (Chothia et al., 1981; Walters and DeGrado, 2006) for these three lipid compositions as well as for the PM composition. This revealed three dimer populations (**Fig. 3A, right panel**): two main right-handed populations with average crossing angles of approximately −27° (RH1) and −9° (RH2) and one minor left-handed population with an average crossing angle of around +9° (LH). To obtain a more detailed view of the dynamical TM dimer association, we plotted the helix crossing angle against the distance between the two Small-X_3_-Small motifs. This revealed several sub-populations associated with each crossing angle peak (**Fig. 3A, left panel**). Notably, membrane lipid composition appeared to modulate this dynamical equilibrium. The PM as well as the 80% DPPC + 20% CHOL compositions favored a RH1 population with a distance between motifs of 6.5 Å. The DPPC membrane allowed a larger diversity of dimer configurations with a slight preference for two types of RH2 populations, with motif distances around 6.5 Å and 8 Å, but also broad RH1 and LH populations. The POPC membrane appeared to depict crossing-angle populations equivalent to PM and 80% DPPC + 20% CHOL compositions but with a shift towards larger motif distances (between 7.5 and 8 Å). Previous studies have shown a fine mechanistic balance between juxtamembrane regions and TM domains (Arkhipov et al., 2013; Defour et al., 2013; Tamagaki et al., 2014). To evaluate the effect of the juxtamembrane (JM) regions on the dynamics of the TM dimer, we performed CG-MD simulations on the TM_24_ segment (**Fig. 1B**), for the three membrane compositions (**Sup. Fig. 4**). Removing the JM regions seemed to shift TM populations towards smaller motif distances (**Sup. Fig. 5**).

**Fig. 3:**
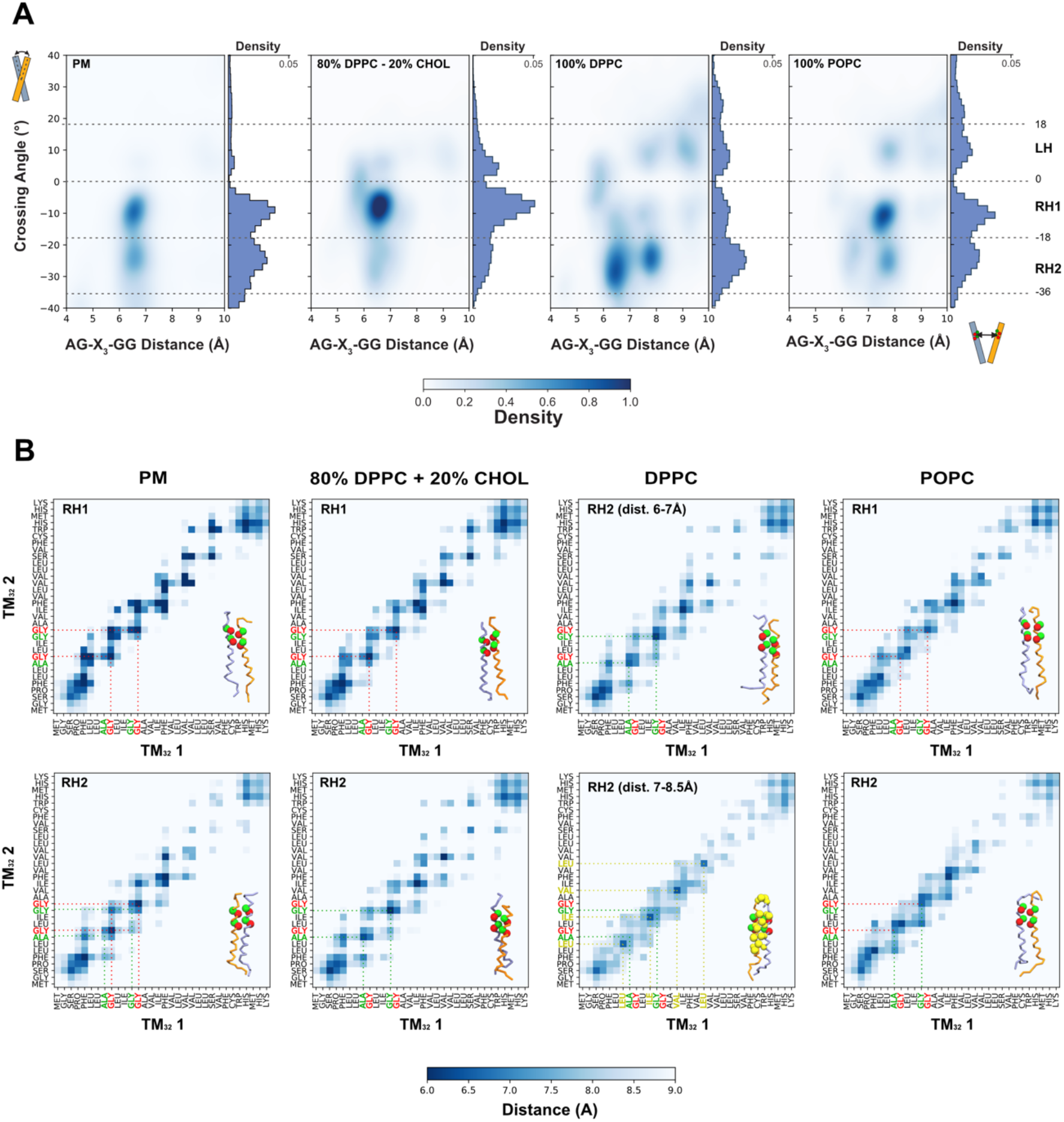
TM dimer dynamics modulated by membrane composition. **A-** TM_32_ helix dimer structural populations for different membrane compositions. A positive value for the crossing angle corresponds to a left-handed (LH) dimer, and a negative value to a right-handed (RH) dimer (details for TM_24_ systems are presented in Sup. Fig. 5). The CG-MD simulations have highlighted two Right Handed conformations (RH1 and RH2) and one Left Handed (LH). RH1 is defined with a crossing angle between −18° and 0°, RH2 between −36° and −18°, and LH between 0° and 18°. **B-** Averaged TM contact matrix extracted from simulations of TM_32_ for the main crossing angle structure in each membrane composition. This highlight the main TM dimerizations via the A_544_-G_548_ motif, G_545_-G_549_ motif or a combination of both motifs.

For each membrane composition, we examined dimer interfaces associated with the different crossing angle populations (**Fig. 3B**). These analyzes showed different TM interactions driven by the interactions of the two Small-X_3_-Small motifs. The interactions through the G_545_-X_3_-G_549_ motif were mostly found in RH1 populations while the A_544_-X_3_-G_548_ motif associations were often related to RH2 populations. In some cases, both motifs interacted together in RH2 (**Fig. 3B**) or LH populations (**Sup. Fig. 6**). We noticed only few events for which the two Small-X_3_-Small motifs were not involved (e. g. second RH2 population for POPC membrane in **Fig. 3B**). We then refined the three main TM configurations seen in our CG-MD simulations (one interaction via A_544_-X_3_-G_548_ motif, one interaction via G_545_-X_3_-G_549_, and one interaction involving both motifs) by performing 400 ns of atomistic MD simulations (see Methods) (**Sup. Fig. 7**). We converted TM dimer structures extracted from the DPPC+CHOL membrane because the atomistic models of the lipids constituting the PM membrane were, to our knowledge, not all parametrized for atomistic force fields. Furthermore, compared to DPPC and POPC membranes, the TM dimer populations in the DPPC+CHOL membrane are the most similar to those in the PM membrane (**Fig. 3A**). For all three structures, the interactions between the Small-X_3_-Small motifs were stable throughout the simulation (**Sup. Fig. 7-B**). Interestingly, the TM dimer involving both motifs seemed more dynamic in term of crossing angle than the two other TM structures (**Sup. Fig. 7-B**).

Thus, MD simulations revealed a dynamic equilibrium of dimer structures involving the two consecutives Small-X_3_-Small motifs, A_544_-X_3_-G_548_ and G_545_-X_3_-G_549_, which may be modulated by membrane composition.

### Distinctive mutations in the Small-X_3_-Small motifs selectively modulate FLRT2 TM dimerization

To assess the individual contributions of the two Small-X_3_-Small motifs to the dimerization, we performed CG-MD simulations with several mutants replacing glycine residues with isoleucine or valine residues, the larger hydrophobic side chains of which are expected to disturb the TM dimerization (Berger et al., 2010; Endres et al., 2013; Heukers et al., 2013) (**Fig. 4A and Sup. Fig. 8**). For each mutant, we evaluated the spatial distributions of the TM_32_ construct embedded in the PM bilayer (**Fig. 4B)**.

**Fig. 4:**
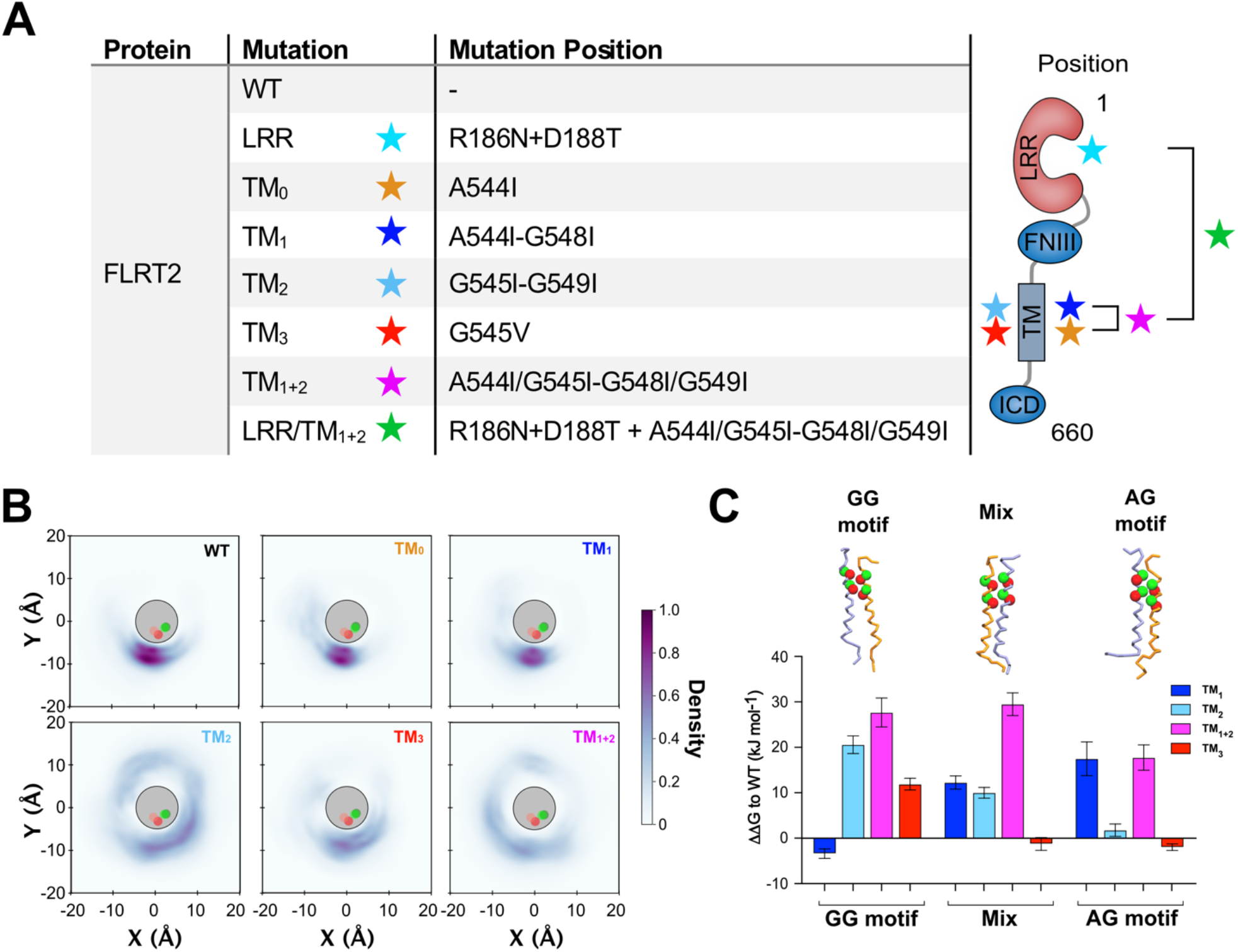
*In silico* mutations in the two Small-X_3_-Small motifs affect the TM dimerization and dynamic equilibrium. **A-** Table of mutations for *in silico* and SMT experiments. The LRR/TM_1+2_ mutant was only used for the SMT experiments. **B-** Spatial distribution profiles of one TM_32_ helix relative to the other for the CG simulations of both WT and mutants in the plasma membrane. The diagram shows the probability density of finding the backbone particles of one TM_32_ helix at a given point in the bilayer plane around the other helix. Green (respectively red) circles depict averaged positions of A_544_ and G_548_ (respectively G_545_ and G_549_) residues. **C-** FEP data showing the effect of the different mutations on the dimer stability. Higher ΔΔG values indicate a more destabilizing mutation effect (more details in Method and Sup. Fig. 10).

Mutations in the A_544_-X_3_-G_548_ motif (mutants TM_0_ and TM_1_) favored formation of a dimer with a spatial distribution focused on the G_545_-X_3_-G_549_ motif while mutations in the G_545_-X_3_-G_549_ motif (mutants TM_2_ and TM_3_) drove interactions through the G_544_-X_3_-G_548_ motif allowing TM domains to explore a wider area. Mutations of both motifs (mutant TM_1+2_) enabled one TM domain to explore the entire bilayer plane surrounding its TM partner, thereby abolishing the specificity of the TM helix interactions. We also performed analyses of the helix crossing angle against the distance between the two Small-X_3_-Small motifs and compared these with the WT distribution (**Sup. Fig. 9-B**). Mutations clearly affected the TM structure populations exploring conformations not seen in the PM membrane but visible in other types of membrane such as DPPC and POPC (**Sup. Fig. 9-B and Fig. 3A**). For the double mutant, the crossing angle density was clearly more diffuse than for the WT or the other mutants, further highlighting a loss of specificity (**Sup. Fig. 9-B**). We then performed these mutations for TM domains embedded in a DPPC bilayer (**Sup. Fig. 2**). For the WT, TM dimer dynamics were clearly different in DPPC than in the PM (**Fig. 3A**). Conversely, the mutants behaved similarly in DPPC and in the PM bilayer, both in term of spatial distribution and crossing angle populations (**Sup. Fig. 9**). Thus, mutants did not seem to be affected by membrane composition.

To further quantify the effect of the mutations on the TM dimerization, we performed non-equilibrium Free Energy Perturbation (FEP) calculations (see Methods). Here, selected residues are perturbed between the WT and mutant states, and the free energy of this change was computed (△G_mut_). By making this change in the context of the dimer or monomer, we can calculate a △△G which quantifies how the mutations affect the relative stability of the dimer (**Sup. Fig. 10B**). The approach of using CG FEP to model mutational △△G has recently been applied in the context of measuring protein-lipid interactions of integral membrane proteins (Corey et al., 2019; Duncan et al., 2020). As we assume that the effect of the mutations on the dimer state might manifest over longer timescales than for lipid interactions, we chose to apply a non-equilibrium protocol (see Methods and **Sup. Fig. 10A**), which allowed us to maximize the sampling of the mutant and WT states. This approach has previously been applied to protein stability studies (Gapsys et al., 2016), as well as to modelling ligand-protein interactions (Gapsys et al., 2021). We performed FEP calculations for WT to TM_1_, TM_2_ and TM_1+2_. These were run using poses with each of the three main dimer interactions: via the A_544_-X_3_-G_548_ motif, via the G_545_-X_3_-G_549_, or through a mix of both motifs (**Fig. 4C**). Each pose was embedded in an 80%DPPC-20%CHOL membrane, which was chosen to keep the membrane as simple as possible for optimal FEP convergence, whilst also recreating dimerization dynamics seen in the PM membrane (**Fig. 3A**). Whilst the TM_1_ mutant impacted mostly TM interactions through the A_544_-X_3_-G_548_ motif and, respectively, the TM_2_ mutant mainly affected TM interactions via the G_545_-X_3_-G_549_ motif, these two mutations only partially disturbed TM dimerization involving both motifs. To control our approach, we tested the TM_3_ (G_545_V). This mutant only moderately disturbed the TM dimers interacting through the G_545_-X_3_-G_549_ motif and did not affect the dimers implicating A_544_-X_3_-G_548_ motif. On the contrary, the double mutant TM_1+2_ strongly destabilized the three poses, with △△G values from around 20 kJ mol^-1^ to 30 kJ mol^-1^. Assuming these mutants near fully destabilize the dimer (as suggested by **Fig. 3C**), this suggests that FLRT2 has a dimerization energy of around 25-30 kJ mol^-1^, similar to estimates for other TM dimers such as those of glycophorin A (Domański et al., 2017; Souza et al., 2021) and ErbA1 (Souza et al., 2021).

Thus, these mutations highlighted two distinct dynamical behaviors of the TM dimer associated with each motif. FEP quantification of TM interactions revealed that only mutation of both motifs together resulted in a ΔΔG value large enough to abolish TM dimerization.

### Mutations in the Small-X_3_-Small motifs affect FLRT2 co-localization in cells

To support the *in silico* results, we performed SMT experiments to assess the contribution of the predicted key residues in the Small-X_3_-Small motifs to dimer formation by mutating the relevant glycine residues to isoleucine or valine (**Fig. 4A**). We tracked FLRT2 receptors on live cells with a sub-pixel accuracy by SMT in two different channels using the dyes Alexa549 and CF640R (**Fig. 5A-C**). Based on receptor frame-to-frame proximity in each channel (**Fig. 5C**), we then built a distribution of the durations of co-localization events (**Fig. 5D**), referred to as t_on_. The duration of co-localization events is a characteristic of the stability of any interaction or association between the tracked receptors, and is independent of expressed receptor concentration (Zanetti-Domingues et al., 2018). Comparison of the t_on_ distributions of WT and mutants (**Fig. 5E,F**) revealed that mutations in only one of the two motifs (either TM_1_, TM_2_ or TM_3_ alone) were insufficient to significantly reduce the baseline average t_on_ of wild-type FLRT2. However, mutation of both Small-X_3_-Small motifs (TM_1+2_ mutant) resulted in a significant shift in the t_on_ distribution towards lower values (**Fig. 5E**). The results are in line with our *in silico* results demonstrating that the two Small-X_3_-Small motifs are required for FLRT interactions *in cis* (**Fig. 4B,C**). These results are also consistent with a previous study showing that mutation of both Small-X_3_-Small transmembrane motifs is necessary to disrupt the EGFR TM dimer and affect receptor function (Endres et al., 2013). In addition to the mutation of both Small-X_3_-Small motifs, a significant shift in t_on_ was also observed for the mutation in the LRR ectodomain, known to abolish FLRT-FLRT trans-interactions (Seiradake et al., 2014), and for the triple mutation LRR+TM_1+2_. In line with the t_on_ results, only diffusion values for the mutants TM_1+2_, LRR, and LRR+TM_1+2_, increased significantly from the WT (**Sup. Fig. 11**). Although the spatial resolution of single molecule tracking is insufficient to discriminate between direct pairwise interactions and co-confinement or joint interactions with the same larger protein complex, this correlation between the decreased duration of co-localization events and the increased diffusion constant of the tracked receptors is consistent with the mutations disrupting interactions that usually occur between WT FLRT2 TM domains.

**Fig. 5:**
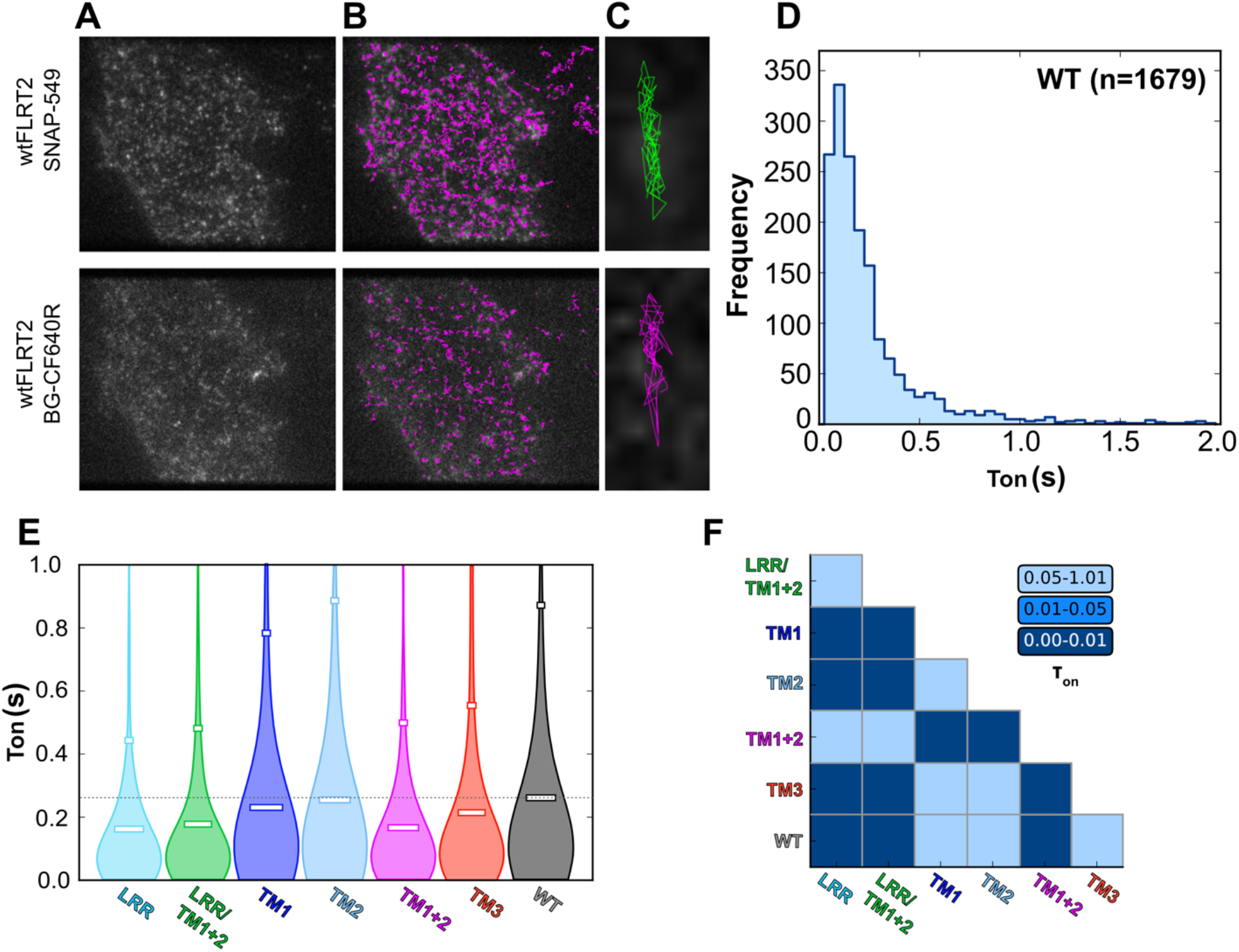
Mutations in the TM domains affect colocalization of FLRT2 monomers at the cell surface. **A-** Single molecule TIRF image of HeLa cells expressing wtFLRT2 labelled with both SNAP-549 and BG-CF640R. **B-** Single molecule tracks are generated from time series of the molecules under observation. **C-** An example pair of colocalized tracks where the tracks are separated by less than 1 pixel (160 nm) during at least 5 frames (250 ms). **D-** Example t_on_ distribution for the WT (n represents the number of tracks analyzed). **E-** Distributions of t_on_ for wtFLRT2 and each of the six FLRT2 mutants tested. **F-** Significance analysis of these distributions based on a Kolmogorov-Smirnov test (more details in Methods section).

Taken together, these data indicate that G_544_-X_3_-G_548_ and G_545_-X_3_-G_549_ motifs in the transmembrane region can both sustain FLRT-FLRT association *in cis*, and that at least one of these motifs is required for wild-type FLRT2 homotypic interactions *in cis*. Interestingly, the LRR ectodomain, which mediates *in trans* FLRT-FLRT interactions (Seiradake et al., 2014), is also required for *in cis* interactions.

## Discussion

Receptor TM dimer association is often a dynamic process involving multiple states and weak interactions, hence direct structural studies remain challenging. As a consequence, only a limited number of TM dimer structures are known and these are often restricted to one state of the TM dimer (Bugge et al., 2016). To make way with understanding TM functions, MD simulations are a method of choice to provide detailed insights into the structures of such assemblies. Here, we have used MD simulations to gain structural insights into the formation of FLRT2 TM dimers. Our models revealed a dynamic equilibrium between conformations involving two successive Small-X_3_-Small motifs, G_544_-X_3_-G_548_ and G_545_-X_3_-G_549_ motifs (**Fig. 1 and 2A**) within a complex lipid bilayer. Our simulations also revealed interactions between the TM domain of FLRT2 and specific lipids (cholesterol, PIP2, PS, and the unsaturated lipids PAPC and DIPE) (**Fig. 2B**). Receptor-lipid interactions are an emerging theme in many signalling systems (Corradi et al., 2018; Duncan et al., 2020) and can affect TM dimerization (Dominguez et al., 2016; Hong and Bowie, 2011; Pawar and Sengupta, 2021). Interestingly, we found that changing the membrane composition modulates the dynamics of FLRT2 TM dimerisation (**Fig. 3**) as do mutations in the Small-X_3_-Small dimerization motifs (**Fig. 4B and Sup. Fig. 9**). As shown by both SMT and MD, targeting both motifs is necessary to significantly affects dimerization (**Fig. 4B,C** and **Fig. 5**).

The TM helices of other receptors, such as EGFRs and EphAs, dimerize via Small-X_3_-Small motifs to transmit extracellular signals to their intracellular enzymatic domains (Bocharov et al., 2010; Endres et al., 2013; Fleishman et al., 2002). There is no enzymatic activity associated with FLRT, which is best known for its functions as a key adaptor protein that defines the structures/functions cell surface signaling hubs (Jackson et al., 2016; Seiradake et al., 2014; Toro et al., 2020), and as a regulator of receptor trafficking (Haines et al., 2006; Leyva-Díaz et al., 2014; Wheldon et al., 2010). Interestingly, dimerization of the EGFR Small-X_3_-Small motif also regulates EGFR trafficking (Heukers et al., 2013) suggesting that *in cis* dimerization via the Small-X_3_-Small motifs may be a conserved feature in the regulation of receptor localization and trafficking, found also in FLRTs. Indeed, the Small-X_3_-Small motifs are conserved in all three FLRT human homologues (FLRT 1-3) and in different species (**Fig. 6A**). Interestingly, the COSMIC database (Forbes et al., 2011) lists a number of cancer-related mutations targeting the TM domain of FLRT2. Two such mutations (A_544_V and G_545_V) map to the Small-X_3_-Small motifs described here, and may affect FLRT2 function and dynamics as seen in MD simulations (**Fig. 3**).

**Fig. 6:**
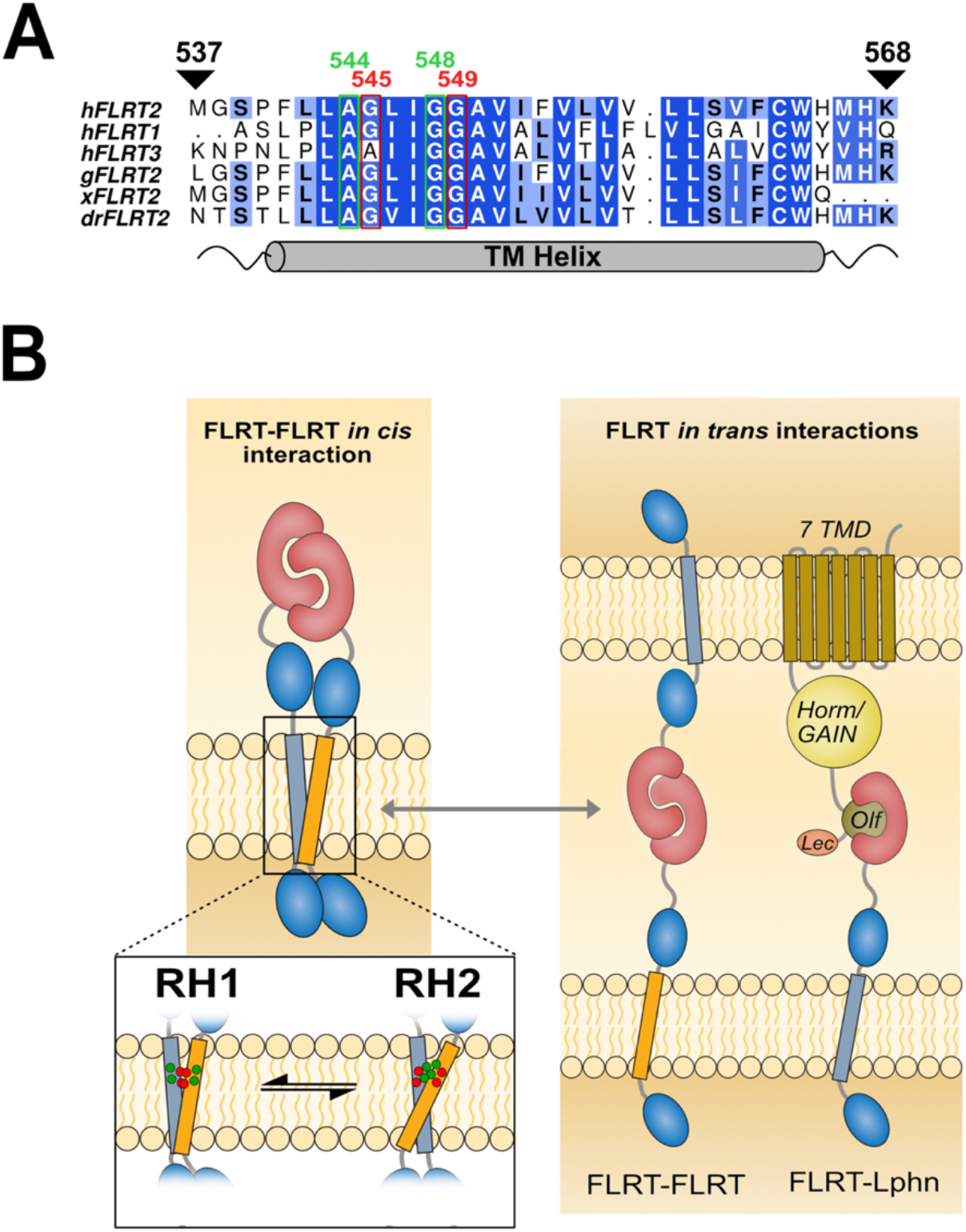
Model of the FLRT *cis*-interaction. **A-** Sequence alignment of the TM domain for FLRT1-3 in human and for FLRT2 in other species (human:h, chicken:g, frog:x, fish:dr). **B-** Model of FLRT2 *cis*-interactions that may compete with different FLRT2 *trans* interactions. The interconversion in between RH1, involving the G_545_-G_549_ motif (in red), and RH2 interactions, driven by the A_544_-G_548_ motif (in green), may be modulated by mutations in the TM domain or environmental conditions such as changes in the lipid composition of the membrane.

Unexpectedly, our results show that the same mutation in the LRR domain that disrupts FLRT-FLRT interactions *in trans* (Seiradake et al., 2014) also disrupts FLRT-FLRT interaction *in cis*, posing the question whether FLRT *cis* and *trans* interactions are competitive. Adding complexity to this issue is the observation that the same mutation also abolishes *trans* FLRT-Lphn interactions (Jackson et al., 2015; Seiradake et al., 2014). These findings suggest that Lphn may also compete with *in cis* FLRT-FLRT dimerization, leading to a mechanism in which FLRTs switch between *in cis* dimerization and different *in trans* interactions via the LRR domain (**Fig. 5B**). Interplay between *cis* and *trans* interactions are key features of typical adhesion proteins, such as cadherins and protocadherins, and is required for effective cell-cell recognition (Honig and Shapiro, 2020). Like other adhesion molecules, FLRTs are broadly expressed. The conformational versatility of its TM domain, and resulting in *cis* binding capability, help explain how these proteins regulate a vast diversity of fundamental developmental processes.

## Materials and Methods

### Modeling Transmembrane domain and Molecular Dynamics Simulations

Results from the PSIpred (Jones, 1999), PRED-TMR2 (Pasquier and Hamodrakas, 1999), and HMMTOP (Tusnady and Simon, 2001) servers were combined to predict the membrane embedded helical region of FLRT2. Twenty-four residues of human FLRT2 (residues 541 – 564) were selected to form the core of the TM helix (TM_24_). The transmembrane domain was created using the Pymol secondary structure creation script: build_seq.py (http://pldserver1.biochem.queensu.ca/~rlc/work/pymol/) and then converted into coarse-grained model. For TM_32_, the four residues both N- and C-terminal of TM_24_ were modelled as random coils using Modeller 9v9 (Webb and Sali, 2016).

Unbiased coarse-grained MD (CG-MD) simulations were performed using GROMACS 4.6 (www.gromacs.org) (Pronk et al., 2013) and GROMACS 2018 (Abraham et al., 2015) with the MARTINI 2.1 forcefield (Marrink et al., 2007; Monticelli et al., 2008). For symmetric membranes (DPPC, POPC and DPPC+CHOL), the temperature was 323K. Electrostatic interactions were shifted to zero between 0 and 1.2 nm and the Lennard-Jones interactions between 0.9 and 1.2 nm. A Berendsen thermostat in combination with a Berendsen barostat with a coupling constant of 1.0 ps, a compressibility of 5.0 × 10^-6^ bar^-1^, and a reference pressure of 1 bar were used. The integration timestep was 20 fs. Simulations were run for either 1 or 2 μs (**Sup. Table 1**) over twenty to thirty replicates to ensure exhaustive sampling of TM helix dimer structures. For the PM membrane, we have used the CHARMM-GUI website (Qi et al., 2015) to create the system. Temperature was maintained at 310K using the V-rescale thermostat (Bussi et al., 2007). Pressure was set to 1 bar using the Parrinello-Rahman barostat (Parrinello and Rahman, 1981) with a coupling constant of 12 ps and a compressibility value of 3 × 10^-4^ bar^-1^. After minimization and equilibration steps, we ran 2 μs of simulations to let the membrane relax. On the final snapshot, we embedded the TM segments and rerun minimization and equilibration steps. To taking into account that this complex system needs longer timescales to equilibrate than symmetric POPC and DPPC membranes, we ran simulations of 4μs (**Sup. Table 1**) over thirty replicates. The integration timestep was 20 fs.

We then converted the three main representative (**Sup. Fig. 7**) coarse grained structures into atomistic models using the CHARMM-GUI MARTINI to All-atom converter (http://www.charmm-gui.org/?doc=input/converter.martini2all). Atomistic simulations were performed with GROMACS 2018 in combination with the CHARMM36 forcefield (Huang and MacKerell, 2013; Lee et al., 2014) and TIP3P water model. The temperature was held at 310K. A first step of energy minimization was performed using the steepest descent algorithm and was equilibrated with a constant temperature ensemble (canonical ensemble, NVT, 310 K) ensemble for 100 ps, followed by a 100 ps equilibration at constant pressure (isothermal-isobaric, NPT, 1 bar). We then ran 100 ns of equilibration by keeping the protein backbone constrained followed by 400 ns of unrestrained production run. We applied a Nosé-Hoover thermostat (Martyna et al., 1992) on the system, coupled with the Parrinello–Rahman barostat (Parrinello and Rahman, 1981), with a compressibility of 4.5×10^-5^ bar^-1^. Long-range electrostatics were modeled using the Particle-Mesh Ewald method (Essmann et al., 1995). All bonds were treated using the LINCS algorithm (Hess, 2008). The integration time step was 1 fs.

### Simulation analysis

Protein and lipid structures were rendered using VMD (Humphrey et al., 1996). Simulations trajectories were analyzed using a combination of Tcl/VMD and Python scripts. Matplotlib was used to create graphs and images of TMD monomer distances, contact matrices, TMD density rendering, and crossing angles analysis. All the scripts used to perform these analyses are available at: https://github.com/MChavent/FLRT. Distances between the two centers of mass of each TM helix were calculated. Density, TM contacts and crossing angle calculations were performed every nanosecond for the part of the trajectory where a dimer was formed. In Figure 3-A (resp. 4-B), the values were renormalized to take into account both the maximum values and time of interactions to properly compare the different membrane (resp. Wild Type and mutants) systems.

### Non-equilibrium free energy perturbation (FEP) calculations

Protein coordinates were extracted from the equilibrium simulations data representing key dimer conformations: one interaction via A_544_-X_3_-G_548_ motif, one interaction via the G_545_-X_3_-G_549_, and one interaction involving both motifs (**Fig. 4C**). For each mutation (TM_1_, TM_2_, TM_1+2_ and TM_3_), side chain beads were added based on the backbone (‘BB’) coordinates.

Each pose was built into solvated membranes of 10 × 10 × 10 nm comprising 80% DPPC and 20% cholesterol using the *insane* protocol (Wassenaar et al., 2015b). CG ions were then added to 0.0375 M (roughly equivalent to 0.15M), and the systems were minimized using the steepest descent method. Two rounds of NPT equilibration were run, first 25 ps with 5 fs timesteps, then 1000 ns with 20 fs timesteps. In both cases the protein ‘BB’ beads had 1000 kJ mol^-1^.nm^-2^ *xyz* positional restraints applied. The temperature was set to 323 K using the V-rescale thermostat (Bussi et al., 2007), with semi-isotropic pressure held at 1 atm using the Berendsen barostat.

For each pose and mutant, non-equilibrium FEP was then carried out (Gapsys et al., 2021). State 0 was set to be the mutant, and state 1 set to be WT. For the relevant residue, this involved the conversion of the BB bead type and setting the sidechain beads to dummy atoms with no LJ or Coulombic interactions. For each state, the system was then minimized using steepest descents, and then simulated for 20 × 100 ns using 20 fs timesteps in the NPT ensemble with the V-rescale thermostat at 323 K (Bussi et al., 2007), and with semi-isotropic pressure held at 1 atm using the Parrinello-Rahman barostat (Parrinello and Rahman, 1981).

For each 100 ns simulation, snapshots were taken every 1 ns from 25-100 ns. Each snapshot was then subjected to 200 ps non-equilibrium FEP (summarized in **Sup. Fig. 10A**). Soft-core potentials on both LJ and Coulombic terms, with an alpha of 0.3, a sigma of 0.25 and a soft-core power of 1. 200 ps calculations were run for the monomer states, which was sufficient for convergence. For the dimer states, 1 ns FEP calculations were run for the TM_1_ mutants, and 4 ns FEP calculations were run for the TM_2_ and TM_1+2_. For the TM_3_ mutations, 200 ps was sufficient sampling for convergence. FEP calculations were run in both the forward (from state 0 to state 1) and backward (from state 1 to state 0) direction. The △G values were then be obtained from the overlap of forward and backward work distributions using the Crooks Fluctuation Theorem (Crooks, 1999). Analyses were carried out using pmx (Gapsys et al., 2015).

Once energies were calculated for each pose with each mutation, △△G values were obtained from the thermodynamic cycle in (**Sup. Fig. 10B**), using the following equation:

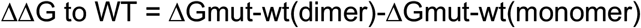

Note that the values for △Gmut-wt(monomer) were obtained by doubling the monomer FEP calculations to account for there being only 1 copy of the FLRT2 TM domain present.

Convergence was tested using 2 metrics. Firstly, consistent variance in FEP values from snapshots taken over the 25-100 ns timescale (**Sup. Fig. 10C**). Second, Convergence analysis measuring the degree of overlap between the forward and reverse FEP calculations (**Sup. Fig. 10D**).

### Cloning

SNAP-FLRT2 was cloned into the EcoRI/XhoI restrictrion sites of the pHSec vector (Aricescu et al., 2006). In SNAP-FLRT2 an N-terminal SNAP tag (containing the RPTPσ signal sequence) was fused to murine FLRT2 (residues A35 – T660) via an HA-tag. Mutations were introduced using molecular cloning.

### Cell Culture and Transfection

HeLa cells were seeded onto uncoated 4-well μ-Slides, #1.5 polymer coverslips (Ibidi) at a density of 1.1×105 cells/well in 600 μL phenol red-free DMEM + 10% FBS + 1% L-Gln + 1% NEAA (complete medium). After 24 h, each well was transfected with 2.0 μg plasmid DNA using FuGENE6, according to the manufacturer’s instructions. Cells were maintained at 37 °C, 5% CO2 and were prepared for experiments 12-18 hours post-transfection.

### BG-CF640R Conjugation

CF640R succinimidyl ester (Biotium) was reacted with BG-NH2 (New England Biolabs) to produce the benzylguanine functionalised dye BG-CF640R. 1 μmol of CF640R succinimidyle ester was reconstituted in DMSO and dissolved in 10 ml 0.1 M sodium bicarbonate buffer (pH 8.4). 1.5 μmol BG-NH2 in DMSO was added to the dye mixture and vortexed well. The reaction was shaken at room temperature overnight before dilution with deionised water. For all subsequent dilutions the conjugation efficiency was assumed to be 100%.

### Two-Colour Fluorescent Labelling

To achieve an approximately equal ratio of single molecules labelled with SNAP Dy549 and BG-CF640R a two-step staining procedure was used. Firstly, the medium was removed from each well of the 4-well μ-Slides and the cells were washed twice with 300 μL complete medium. BG-CF60R was diluted in complete medium to a final concentration of 10 nM and applied to each well of the μ-Slide for 5 min. The medium was then exchanged for 150 μL 10 nM SNAP-Dy549 (SNAP-Surface 549, New England Biolabs) in complete medium and incubated for a further 5 min. All labelling steps were performed at 37°C, 5% CO2. Labelled cells were then washed three times with complete medium and the final wash replaced with Live Cell Imaging Solution plus 1:50 ProLong Antifade reagent (both ThermoFisher) and incubated for at least 15 min, at 37 °C, 5% CO2 before beginning experiments.

### Single molecule image acquisition and feature tracking

Single-molecule images were acquired using an Axiovert 200M microscope with an iLas2 TIRF illuminator (Cairn, UK), with a ×100 oil-immersion objective (α-Plan-Fluar, NA= 1.46; Zeiss, UK) and an EMCCD (iXon X_3_; Andor, UK). The microscope is also equipped with a wrap-around incubator (Pecon XL S1). The 561 and 642 nm lines of a LightHub laser combiner (Omicron-laserage Laserprodukte GmbH) were used to illuminate the sample and an Optosplit Image Splitter (Cairn Research) was used to separate the image into its spectral components as described previously (Webb et al., 2006). The field of view of each channel for single-molecule imaging was 80 × 30 μm. Typically, for each condition at least 50 fields of view comprising one or more cells were acquired from a total of 4 independent biological replicates. Single molecules were tracked in each field of view for 30s, by which time the majority of molecules had undergone photobleaching. All single-molecule time series data were analyzed using the multidimensional analysis software described previously (Rolfe et al., 2011). Briefly, this software performs frame-by-frame Bayesian segmentation to detect and measure features to sub-pixel precision, then links these features through time to create tracks using a simple proximity-based algorithm. The software determines cubic polynomial registration transformations between wavelength channels from images of fluorescent beads. Feature detection and tracking was performed independently in each channel.

### Calculation of colocalisation and t_ON_

Two-colour TIRF images of the basolateral surfaces of cells were chromatically separated by a beam splitter and registered using custom-made software to map the relative positions of the probes over the time course of data acquisition (Rolfe et al., 2011) and extract single molecule tracks. A colocalisation event was defined as one in which a track in one channel moves within one pixel of a track in the other channel before they move apart again. The duration of each such event is one measurement of t_ON_. This parameter indicates the stability of presumptive receptor interactions while being insensitive to variation in expression of the receptors between cells or different levels of labelling with the two probes within cells (Zanetti-Domingues et al., 2018). The track positions were registered between channels prior to this analysis. To reduce the impact of localisation error on these results a temporal Gaussian smoothing filter of FWHM 4 frames (200 ms) was applied to the position traces before the colocalisation analyses. t_ON_ distributions were compared between conditions using the two-sample Kolmogorov-Smirnov test to decide which were significantly different.

### Mean squared displacement and diffusion calculation

From single particle tracks, mean squared displacement (MSD) curves were calculated as MSD(*ΔT*)=<|*r_i_*(*T*+*ΔT*)-*r_i_*(*T*)|^2^> where |*r_i_*(*T*+*ΔT*)-*r_i_*(*T*)| is the displacement between position of track *i* at time *T* and time *T*+*ΔT* and the average is over all pairs of points separated by *ΔT* in each track. The average instantaneous diffusion coefficient (D) for these tracks was calculated by fitting a straight line to the first two points of the MSD curve then calculating D directly from the gradient m of the fit, D=m/4. The tracks for each single molecule field of view (FOV) were pooled into one MSD curve per FOV to produce a sample of D values, one value per FOV per condition. These D distributions were compared between conditions using the Kolmogorov-Smirnov test to decide which were significantly different. The two-sample KS test is a non-parametric test of the null hypothesis that two independent samples are drawn from the same continuous distribution. We use the 2-sided KS test implemented in Python scipy.stats.ks_2samp function.

## Acknowledgements

A.L.D. is supported by the BBSRC grant BB/R00126X/1 and Pembroke College, Oxford (BTP Fellowship). E.Y.J. was funded by UK Medical Research Council programme grant G9900061. The Wellcome Centre for Human Genetics is supported by Wellcome Trust Centre grant 203141/Z/16/Z. E.S. is funded by the Wellcome Trust (202827/Z/16/Z), EMBO YIP and supported by the COST action ‘Adhere’n Rise’. V.A.J was funded by the Wellcome Trust DPhil programme in Cellullar Structural Biology. M.S.P.S. is funded by Wellcome Trust grants WT092970MA and 208361/Z/17/Z. M. C. is supported by the CNRS-MITI grant “Modélisation du vivant” 2020. This work was granted access to the HPC resources of CALMIP supercomputing center under the allocation 2020-17036. The single molecule analysis used computing resources provided by STFC Scientific Computing Department’s SCARF cluster. We acknowledge Life Science Editors for proofreading the manuscript. M. C. thanks E. Haanappel and A. Atkinson for fruitful discussions and support.

## Authors contributions

Conceptualization: ACK, MSPS, ES, MLMF, MC

Methodology: CTJ, DJR, RAC, AC

Investigation: VJ, CJT, DJR, RAC, ALD, MN, AC, MC

Supervision: ACK, MSPS, MLMF, ES, MC

Writing—original draft: VJ, JH, CJT, DJR, MSPS, MLMF, ES, MC

Writing—review & editing: CJT, DJR, RAC, ALD, ACK, EYJ, MSPS, MLMF, ES, MC

## Data and materials availability

Scripts used to analyze MD simulations and models for the main conformations both in coarse grained and atomistic representations are available at: https://github.com/MChavent/FLRT. Additional data related to this paper may be requested from the authors.

