## supplementary material for "The guidance and adhesion protein FLRT2 dimerizes *in cis* via dual Small-X_3_-Small transmembrane motifs"

**This PDF file includes:**

**Fig. S1:** Plasma membrane composition.
**Fig. S2:** Distance between the $TM_{32}$ monomers in different type of membranes.
**Fig. S3:** Averaged contact matrices for $TM_{24}$ and $TM_{32}$ in different membrane systems.
**Fig. S4:** Distance between the $TM_{24}$ monomers in different type of membranes.
**Fig. S5:** Helix crossing angle distributions for $TM_{24}$ dimers.
**Fig. S6:** Averaged TM contact matrix extracted from simulations of $TM_{32}$ in 80% DPPC and 20% of cholesterol for the LH population.
**Fig. S7:** Refinement of CG models using atomistic simulations.
**Fig. S8:** Distance between the $TM_{32}$ monomers mutants in PM.
**Fig. S9:** Effects of mutations on the dynamics of the TM-dimer.
**Fig. S10:** FEP calculations.
**Fig. S11:** Mutations of the TM domains affect diffusion of FLRT2 receptor.
**Table S1:** Summary of unrestrained simulations.

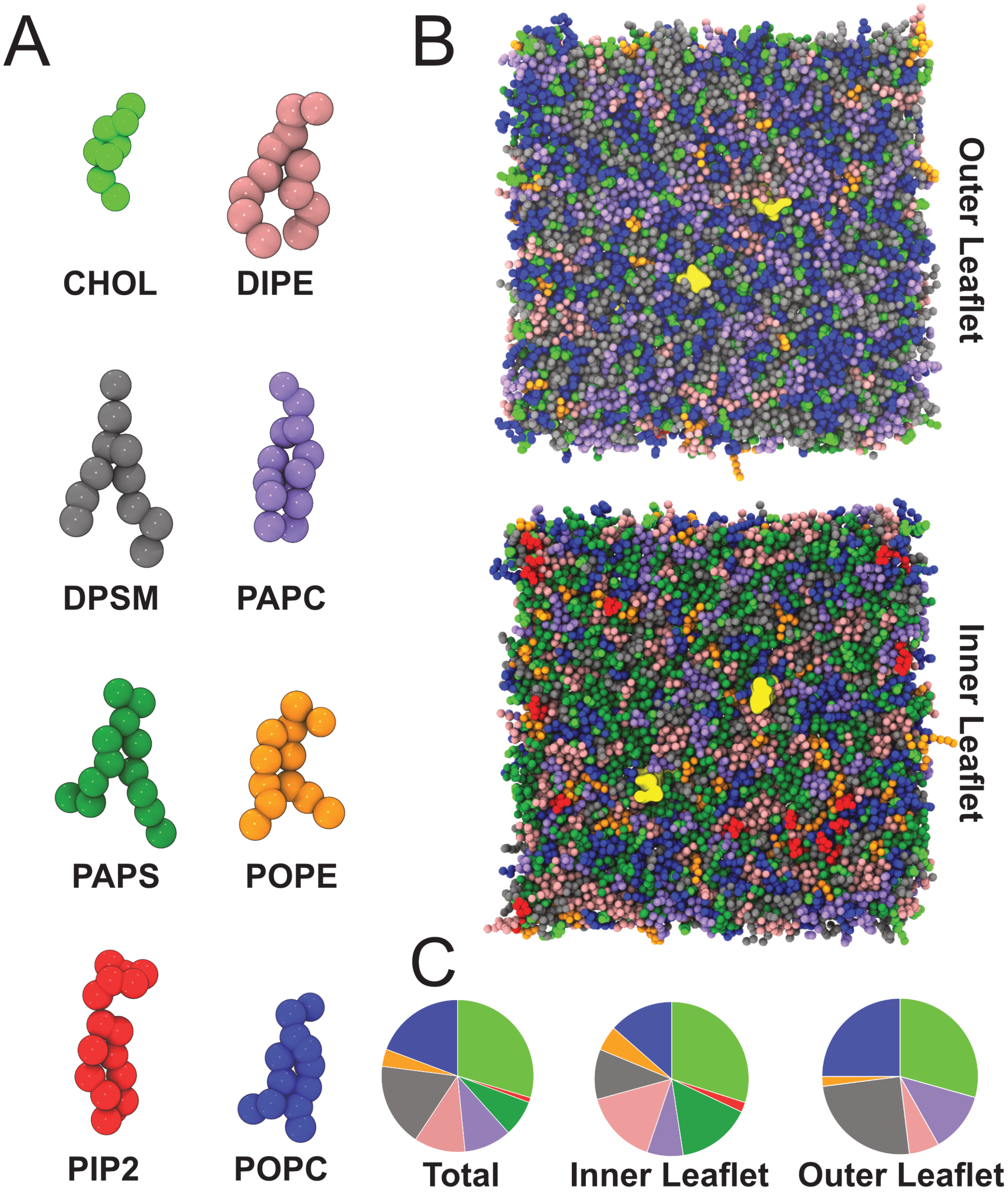


**Fig. S1: Plasma membrane composition. A-** Coarse-grained models of each lipid constituting the plasma membrane. **B-** Equilibrated model of the plasma membrane model with colors corresponding to colored lipids in A. In yellow the starting positions of the two $TM_{32}$ domains. **C-** Lipid distributions for each leaflet and in total.

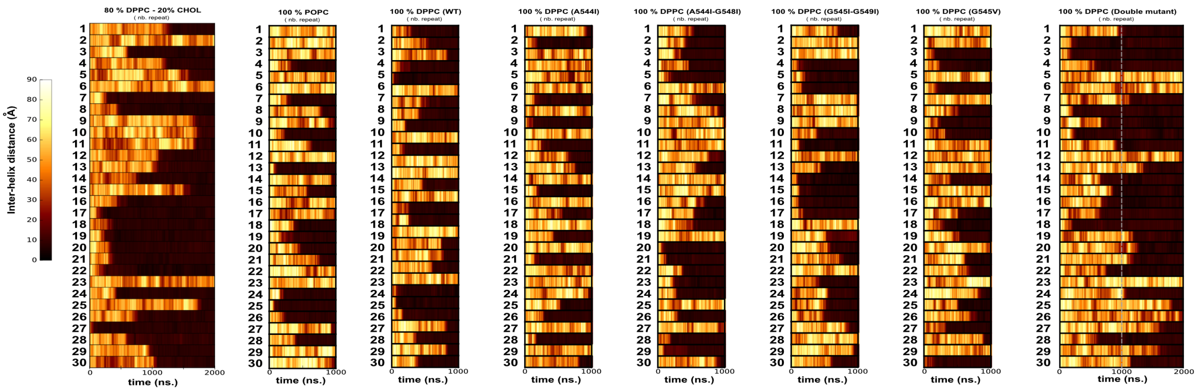


**Fig. S2: Distance between the $TM_{32}$ monomers in different type of membranes.** Distance between the two $TM_{32}$ helices as a function of the time during the course of the CG-MD simulation for the WT and mutants (double mutant corresponds to A544I-G548I+G545I-G549I, see Fig. 4A for the mutations details). The two helices are first positioned 60 Å apart and then freely diffuse in the membrane. For the double mutant, too few simulations depicted interacting domains after 1 µs to perform meaningful analysis, so these simulations were extended to 2 µs.

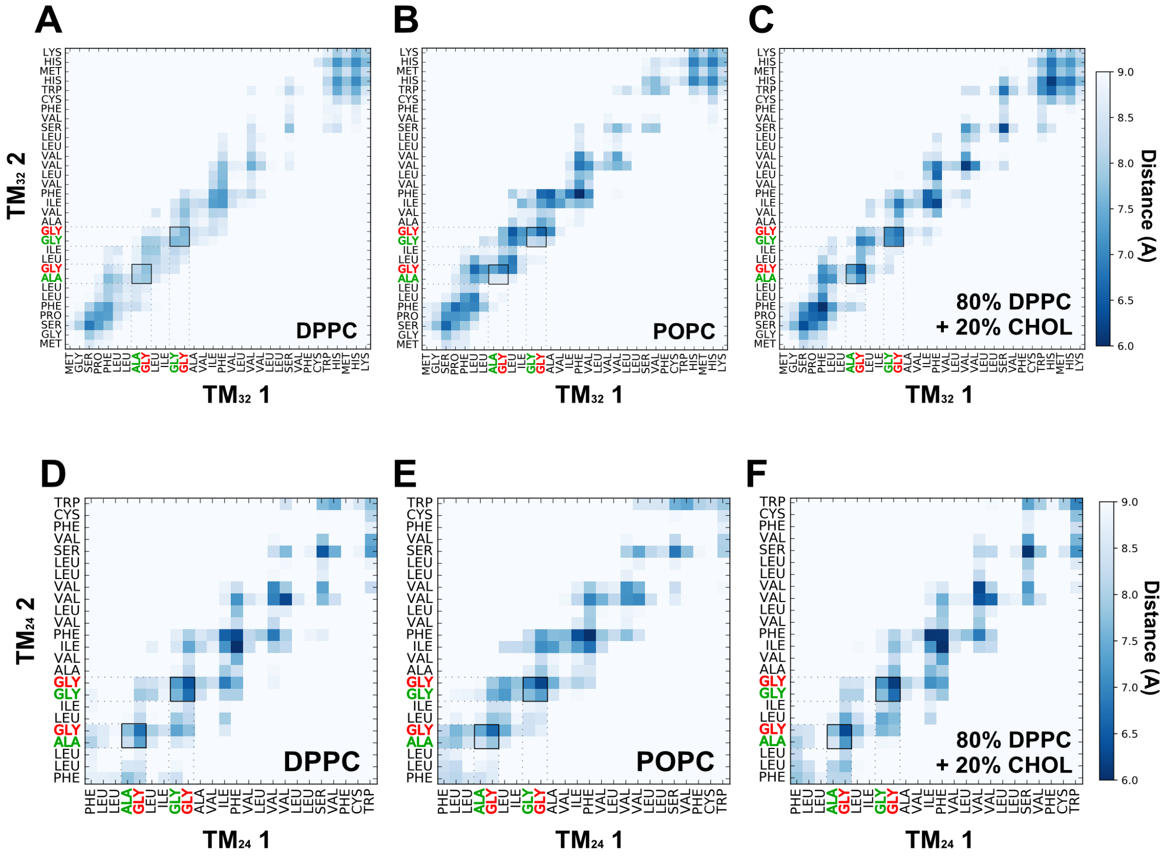


**Fig. S3: Averaged contact matrices for $TM_{24}$ and $TM_{32}$ in different membrane systems.** For each membrane composition, averaged $TM_{32}$ (**A**,**B**,**C**) and $TM_{24}$ (**D**,**E**,**F**) contact matrices were calculated for parts of the simulation when TM domains are in interaction (see Sup. Fig. 1,2).

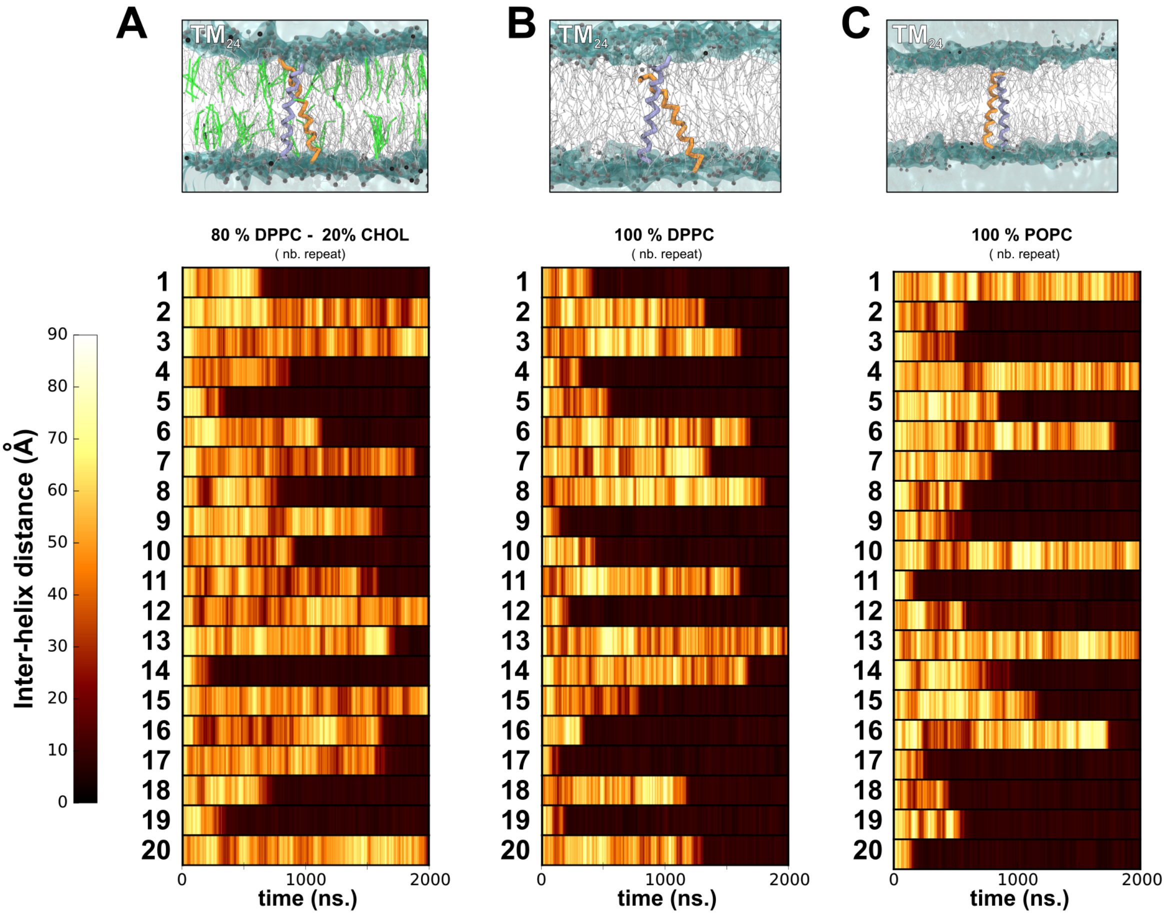


**Fig. S4: Distance between the $TM_{24}$ monomers in different type of membranes.** Distance between the two $TM_{24}$ helices as a function of time during the course of the CG-MD simulation for 3 different membrane compositions: 80% DPPC + 20% cholesterol (**A**), 100% DPPC (**B**), and 100% POPC (**C**). The 2 helices are first positioned 60 Å apart and then freely diffuse in the membrane. Top: one representative depiction of the $TM_{24}$ dimer configuration at the end of the simulation.

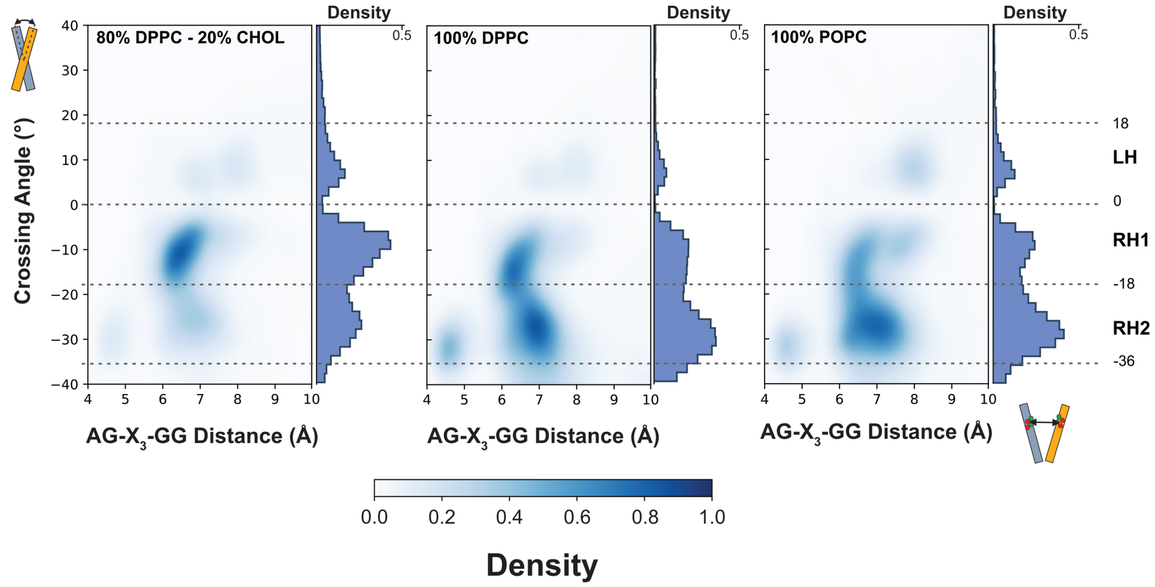


**Fig. S5: Helix crossing angle distributions for TM24 dimers.** Averaged helix crossing angle distribution for TM24 systems based on individual simulations for different membrane compositions in which TM domains interacted for at least two hundreds of nanoseconds (see Sup. Fig. 2).

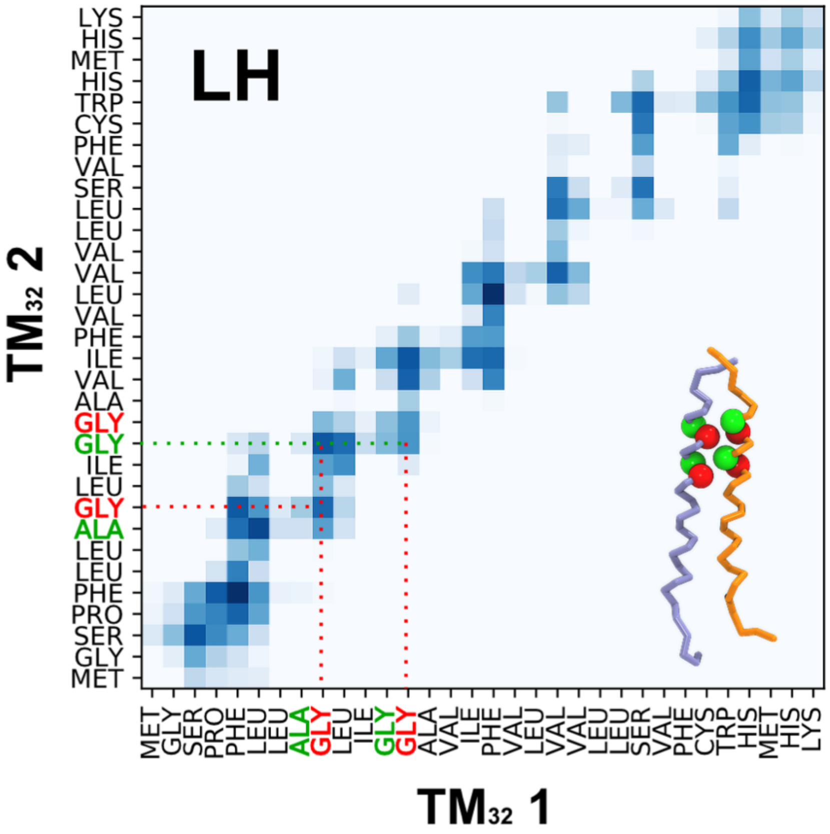


**Fig. S6: Averaged TM contact matrix extracted from simulations of $TM_{32}$ in 80% DPPC and 20% of cholesterol for the LH population** (see Fig. 3A) displaying a TM interaction involving both $G_{544}$-$X_3$-$G_{548}$ (green) and $G_{545}$-$X_3$-$G_{549}$ (red) motifs.

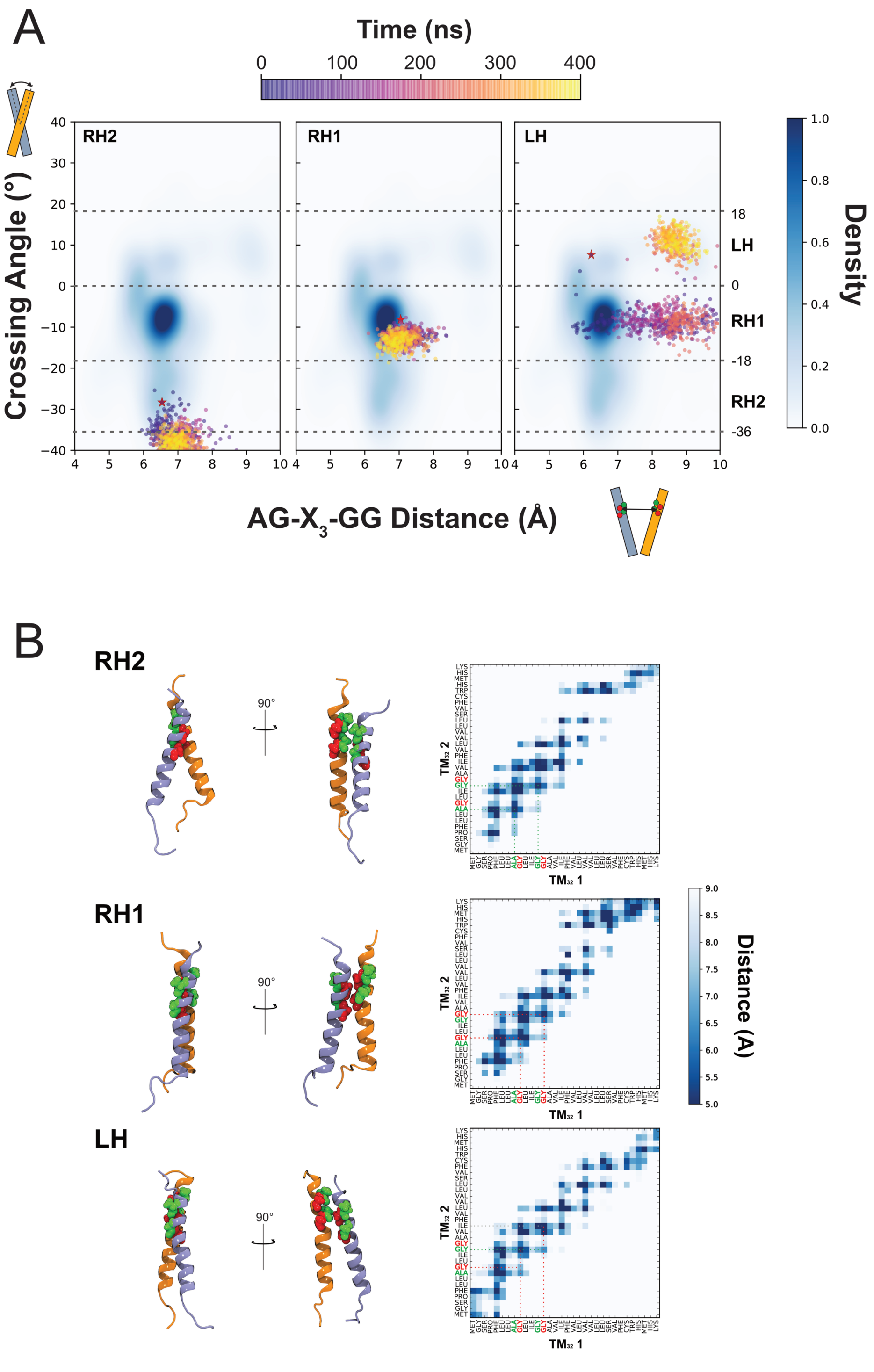


**Fig. S7: Refinement of CG models using atomistic simulations. A-** We converted in atomistic representation the three main representative structures of the dimer embedded in a 80% DPPC + 20% cholesterol membrane. For each representative conformation, crossing angles populations based on atomistic simulations are colored in function of the time. These crossing angles are superimposed to the averaged population obtained from CG simulations. The starting structure is represented by a red star. **B-** In each case, a contact map calculated during the course of the 400 ns simulation and a final structure of the helix dimer are shown.

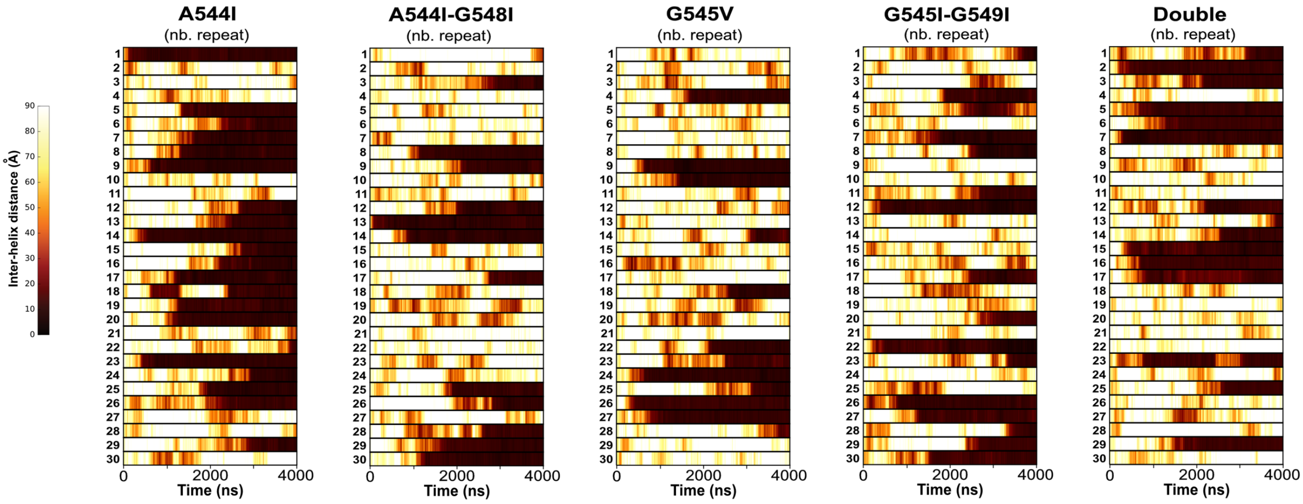


**Fig. S8: Distance between the $TM_{32}$ monomers mutants in PM.** Distance between the two $TM_{32}$ helices as a function of the time during the course of the CG-MD simulation for the mutants in the PM. The two helices are first positioned 60 Å apart and then freely diffuse in the membrane.

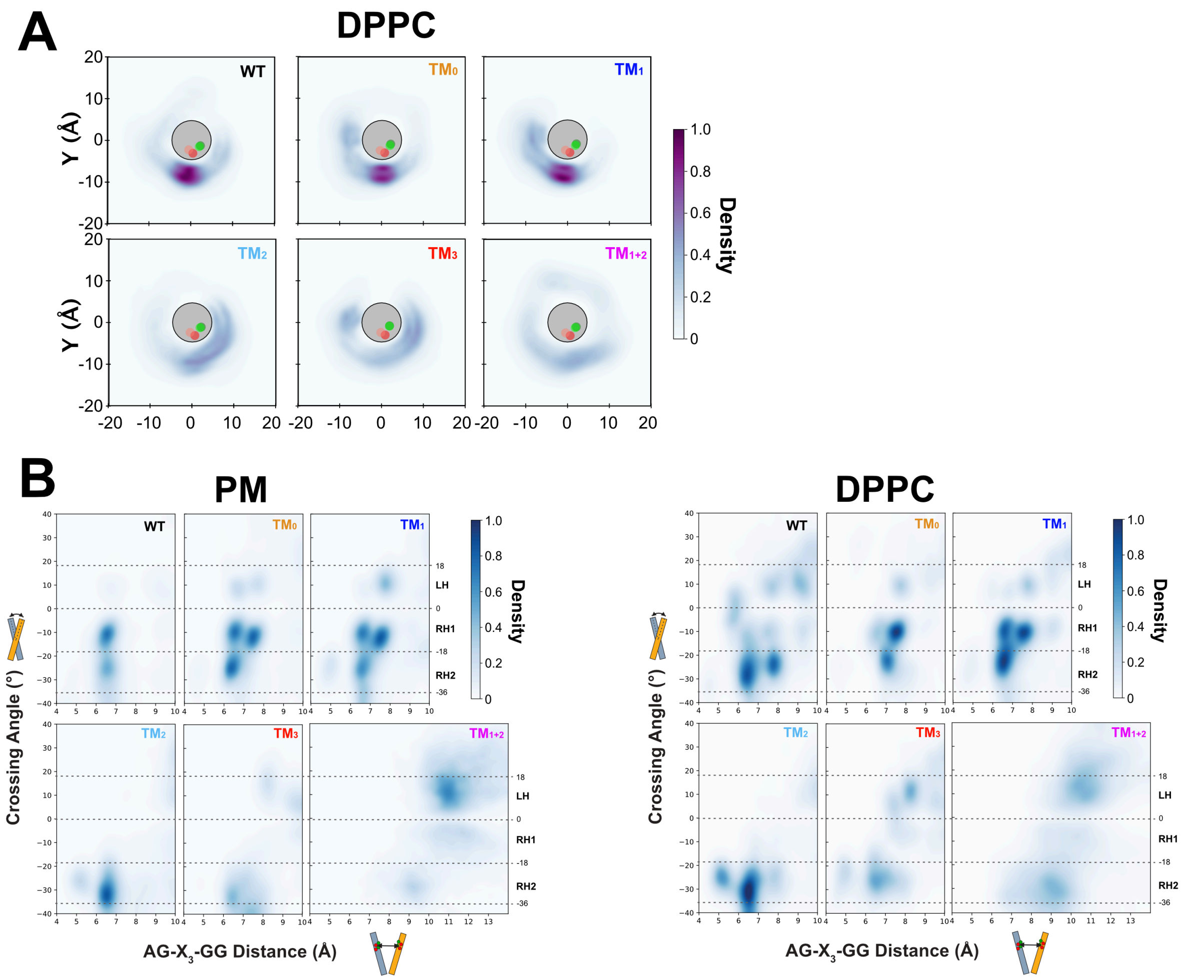


**Fig. S9: Effects of mutations on the dynamics of the TM-dimer. A-** Spatial distribution profiles of one $TM_{32}$ helix relative to the other for the CG simulations of both WT and mutants in a DPPC bilayer. **B-** $TM_{32}$ WT and mutants structural populations for plasma membrane and DPPC bilayer.

**Figure Supp**

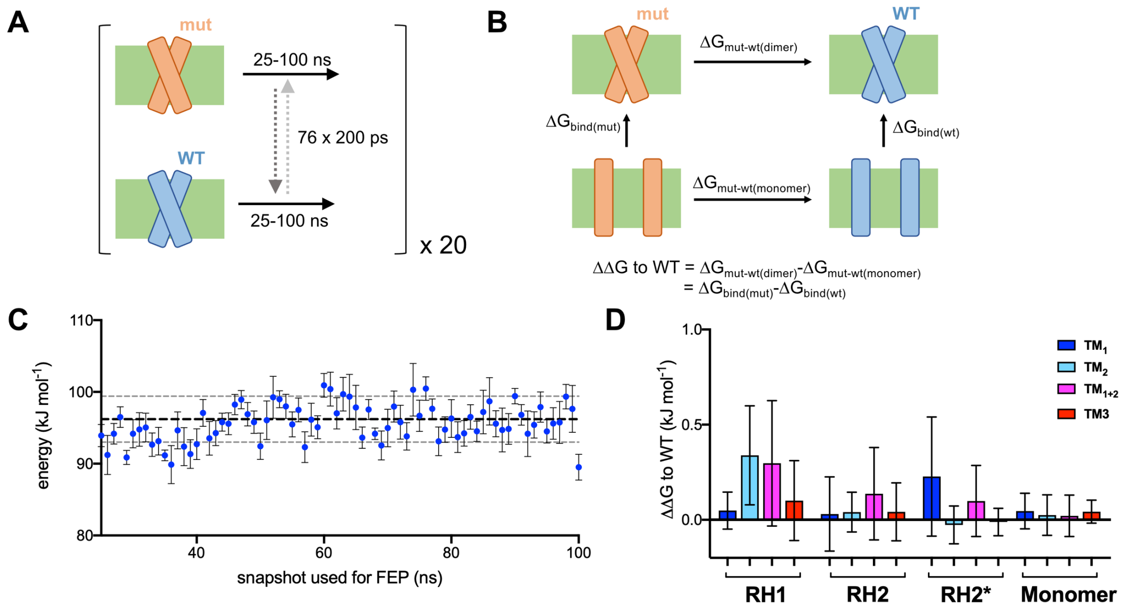


**Fig. S10: FEP calculations. A-** Overview of non-equilibrium FEP scheme. For each dimer/monomer system, 100ns simulations are run in state 0 ('mut') or state 1 ('WT'). Snapshots are taken every 1 ns from 25-100 ns, and subjected to 200 ps FEP from either state 0 to 1 or vice versa. This process is repeated 20 times for statistics. **B-** Thermodynamic cycle for calculating ΔΔG values. The top and bottom lateral ΔG values are explicitly calculated here in the non-equilibrium FEP, allowing calculation of a ΔΔG using the equation. **C-** Example plot showing how the ΔGmut-WT(dimer) values change depending on input snapshot. A regular spread of data across the data suggests that the dimers are stable and are not shifting in conformation during the equilibrium simulation. Shown are the data for the RH2 TM1+2 mutant, which gives one of the higher ΔΔG values so would be expected to change the most. **D-** Convergence analysis measuring the degree of overlap between the forward and reverse FEP calculations, based on (Hahn and Then, 2010) and implemented in pmx (Gapsys et al., 2015). Here, a value of less than ca. 0.8 is considered as having sufficient overlap to show convergence.

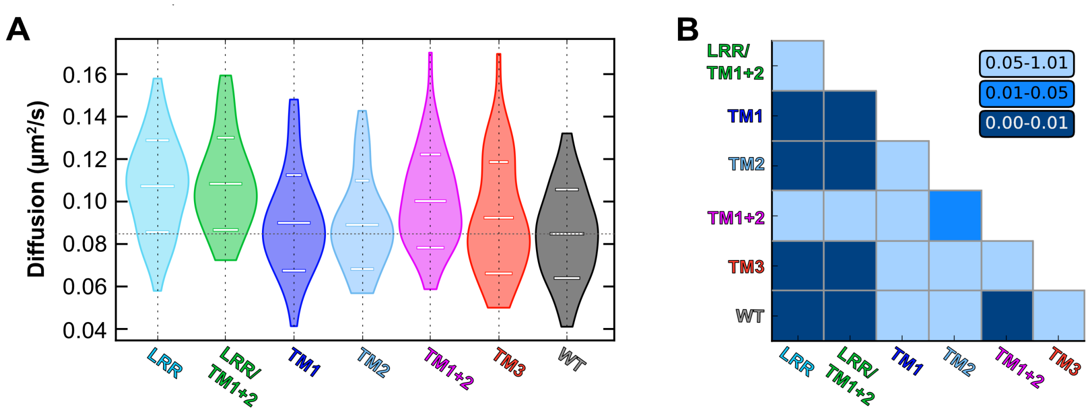


**Fig. S11: Mutations of the TM domains affect diffusion of FLRT2 receptor. A-** Distribution of instantaneous diffusion coefficient (D) for wtFLRT2 and each of the six FLRT2 mutants tested. **B-** Significance analysis of the distributions presented in A based on a Kolmogorov-Smirnov test (more details in Methods section).

**Table S1: Summary of unrestrained simulations.** CG simulations contained from c.a. 10600 particles (POPC, DPPC, and DPPC+CHOL membranes) to c.a. 32500 particles (PM).

| **Coarse-Grain** | | | | |
|---|---|---|---|---|
| Protein | Mutation | Bilayer | Simulation time [μs] | Number of repeats |
| -- | -- | Plasma Membrane | 2 | 1 |
| FLRT2 $TM_{24}$ dimer | WT | 100% POPC | 2 | 20 |
| | | 100% DPPC | 2 | 20 |
| | | 20% CHOL 80% DPPC | 2 | 20 |
| FLRT2 $TM_{32}$ dimer | WT | Plasma Membrane | 4 | 30 |
| | | 100% POPC | 1 | 30 |
| | | 100% DPPC | 1 | 30 |
| | | 20% CHOL 80% DPPC | 2 | 30 |
| | WT (AT) | 20% CHOL 80% DPPC (RH1) | 0.5 (0.1 + 0.4) | 1 |
| | | 20% CHOL 80% DPPC (RH2) | 0.5 (0.1 + 0.4) | 1 |
| | | 20% CHOL 80% DPPC (LH) | 0.5 (0.1 + 0.4) | 1 |
| | A544I ($TM_0$) | Plasma Membrane | 4 | 30 |
| | | 100% DPPC | 1 | 30 |
| | A544I-G548I ($TM_1$) | Plasma Membrane | 4 | 30 |
| | | 100% DPPC | 1 | 30 |
| | G545I-G549I ($TM_2$) | Plasma Membrane | 4 | 30 |
| | | 100% DPPC | 1 | 30 |
| | G545V ($TM_3$) | Plasma Membrane | 4 | 30 |
| | | 100% DPPC | 1 | 30 |
| | A544I-G548I + G545I-G549I ($TM_{1+2}$) | Plasma Membrane | 4 | 30 |
| | | 100% DPPC | 2 | 30 |
